# 3D printing of Microgel-loaded Modular LEGO-like Cages as Instructive Scaffolds for Tissue Engineering

**DOI:** 10.1101/2020.03.02.974204

**Authors:** Christina Hipfinger, Ramesh Subbiah, Anthony Tahayeri, Avathamsa Athirasala, Sivaporn Horsophonphong, Greeshma Thrivikraman, Cristiane Miranda França, Diana Araujo Cunha, Amin Mansoorifar, Albena Zahariev, James M. Jones, Paulo G. Coelho, Lukasz Witek, Hua Xie, Robert E. Guldberg, Luiz E. Bertassoni

**Affiliations:** Division of Biomaterials and Biomechanics, Department of Restorative Dentistry, School of Dentistry, Oregon Health and Science University, Portland, OR, USA; School of Dentistry, Mahidol University, Bangkok, Thailand; Division of Biomaterials and Biomimetics, School of Dentistry, New York University, New York, NY, USA.; Phil and Penny Knight Campus for Accelerating Scientific Impact, University of Oregon, Eugene, OR, USA; Center for Regenerative Medicine, School of Medicine, Oregon Health and Science University, Portland, OR, USA; Department of Biomedical Engineering, School of Medicine, and Cancer Early Detection Advanced Research Center (CEDAR), Knight Cancer Institute, Oregon Health and Science University, Portland, OR, USA

## Abstract

Biomaterial scaffolds have served as the foundation of tissue engineering and regenerative medicine. However, scaffold systems are often difficult to scale in size or shape in order to fit defect-specific dimensions, and thus provide only limited spatiotemporal control of therapeutic delivery and host tissue responses. Here, a lithography-based three-dimensional (3D) printing strategy is used to fabricate a novel miniaturized modular LEGO-like cage scaffold system, which can be assembled and scaled manually with ease. Scalability is based on an intuitive concept of stacking modules, like conventional LEGO blocks, allowing for literally thousands of potential geometric configurations, and without the need for specialized equipment. Moreover, the modular hollow-cage design allows each unit to be loaded with biologic cargo of different compositions, thus enabling controllable and easy patterning of therapeutics within the material in 3D. In summary, the concept of miniaturized cage designs with such straight-forward assembly and scalability, as well as controllable loading properties, is a flexible platform that can be extended to a wide range of materials for improved biological performance.

**TOC:** **3D printed LEGO-like hollow microcages** can be easily assembled, adjoined, and stacked-up to suit the complexity of defect tissues; aid spatial loading of cells and biomolecules; instruct cells migration three-dimensionally; and facilitate cell invasion and neovascularization *in-vivo*, thus accelerating the process of tissue healing and new tissue formation.

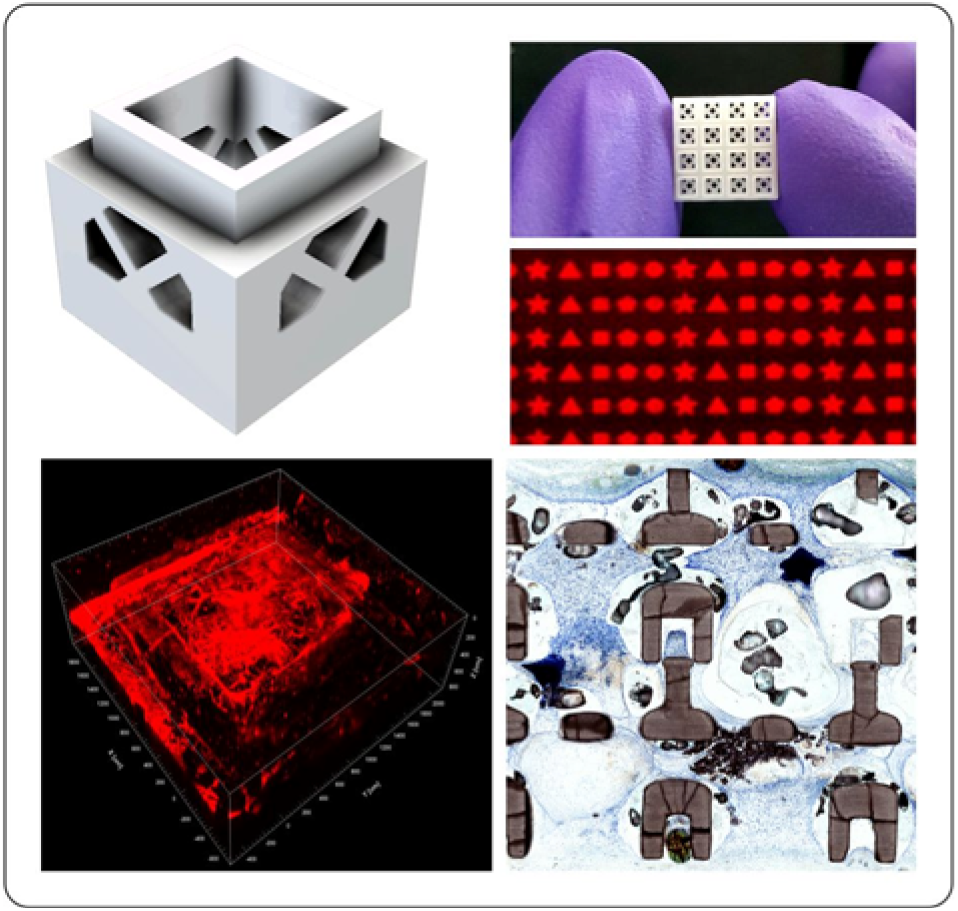

---

Additive manufacturing has facilitated significant progress in scaffold design and fabrication for regenerative medicine applications.(1, 2) These have been especially promising for the engineering of improved scaffold materials for regeneration of large tissue defects, which are intended to be used for patient-specific design parameters, such as defect shape, size and three-dimensional (3D) architectural complexity.(3, 4) Therefore, the fabrication of implantable constructs that can complement size, shape and volumetric complexity of a given defect can benefit from the development of manufacturing strategies that are easily scalable, can be performed entirely in the operating room, and allow for assembly of multiple geometries with ease,(5, 6) without the need for specialized equipment or personnel. An ideal scaffold system would not only be compatible with defect-specific architectures but also with controllable loading of biologics (*i.e.*, cells, growth factors, hydrogels, and combinations thereof), in a simple and site-specific manner,(7) in order to enhance the spatial and temporal control of host tissue ingrowth within the grafted material. Strategies that address these criteria have remained elusive to date. Our team has developed a new scaffold assembly system which takes advantage of 3D-printed scaffolds fabricated in the form of rigid, hollow and stackable miniaturized cage modules, resembling LEGO blocks, which allow for easy and scalable configuration of complex geometries via an intuitive manual assembly process. The ability to design such scaffolds with hollow features, enables loading of cargo with different compositions in controllable arrangements, as to fabricate scaffolds with spatially-defined instructive cues. As a proof of principle, we employed 3D-printed LEGO-like cages loaded with microscale granular hydrogels (microgels)(8) supplemented with growth factors of varying compositions, which show enhanced cell invasion into the core of assembled LEGO-constructs in a controllable manner, thus accelerating the process of new tissue formation and healing.

In order to evaluate the functionality of these 3D-printed modular miniaturized cages, the LEGO-like modules were fabricated using the design configurations that are common to conventional LEGO blocks. Hydrogel-laden LEGO-like scaffolds composed of high-density β-TCP ceramic were 3D printed using a Lithography-based Ceramic Photopolymerization (LCM) method.(9) Similar LEGO-like scaffolds from other rigid polymers can also be fabricated using standard digital light projection (DLP) 3D printers (**Supplementary Figure 1A, 1B**).(10) To that end, repeating units of 3.375 mm^3^ ceramic cages with 1.5×1.5×1.5 mm hollow dimensions, and 230-560 µm thick walls, were fabricated as vertically stackable blocks with repeating units of 1×1 (Figure 1A-C, 1E, Supplementary Figure 1C), 2×2 (Figure 1F), and 4×4 (Figure 1D, 1G) configurations. Blocks were assembled in a multi-stacked fashion with varying dimensions and a consistent outline across the perimeter of the samples, semi-pyramidal shapes with a larger base and a smaller top (Figure 1H and 1J), as well as randomly oriented irregular outlines (Figure 1I). Variations of this stackable design functionality make it possible, for instance, to achieve a total of 29,413 configurations out of only 4 layers of 4×4 blocks, which illustrates the range of possibilities for assembling multi-structured constructs out of individual modules.

**Figure 1.**
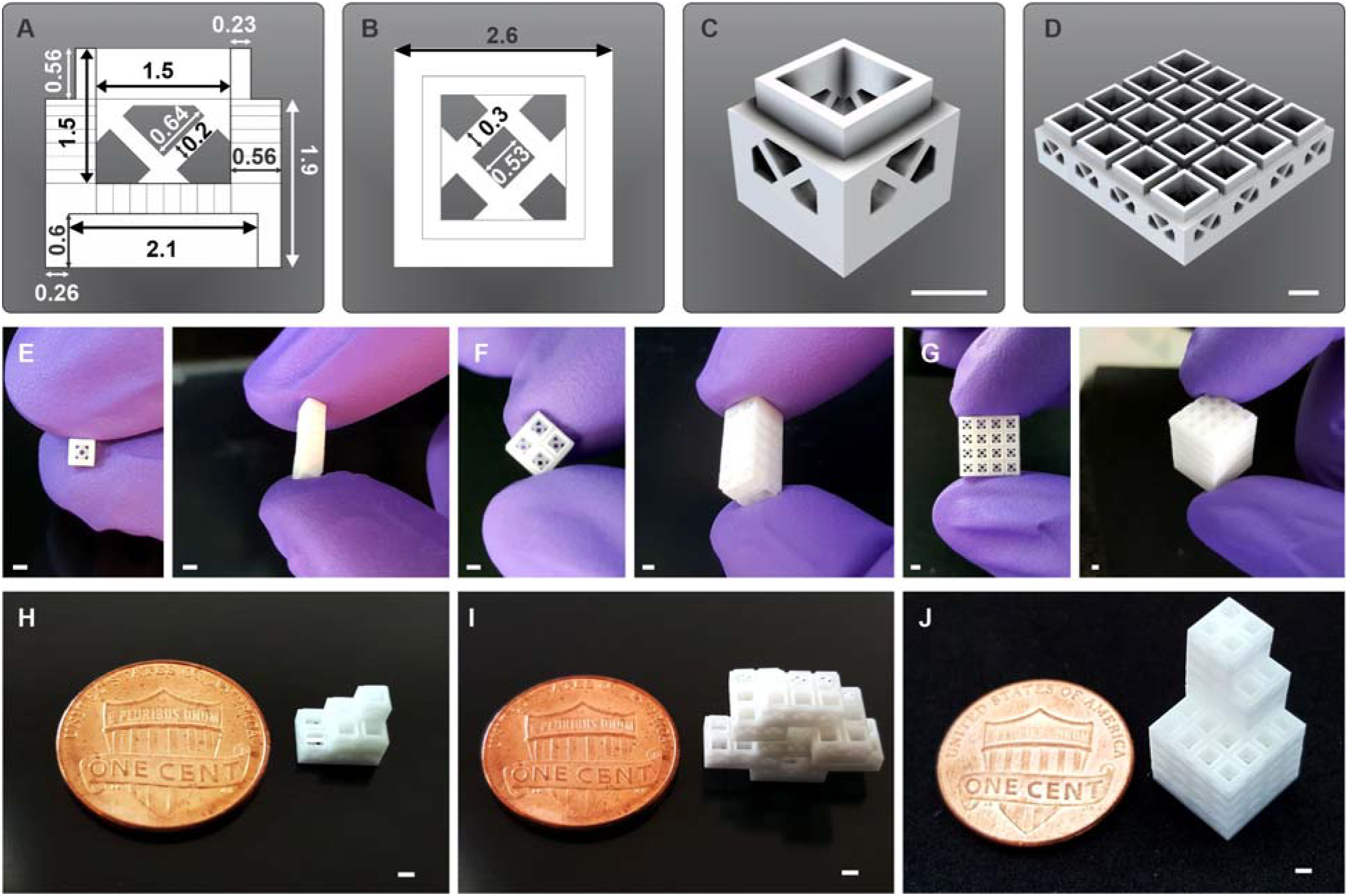
A, B, C) Schematic designs of a 1×1 scaffold from a vertical, horizontal and orthogonal perspective. D) Orthogonal projection of 3D design of a 4×4 LEGO cage scaffold. E) Photographs of single 1×1 (left) and stacked-up LEGO scaffolds (right). F, G) Assembled LEGO scaffolds from 2×2 and 4×4 designs (left). H-J) Photographs of LEGO scaffolds assembled with semi-pyramidal shapes as well as randomly oriented irregular outlines. (scale bar: 1.5 mm)

An additional, key functionality of the hollow cage design is the ability to load this scaffold with cargo in a spatially controllable manner. Strategies to engineer tissue constructs with heterogeneous structure and composition remain scarce,(11) and these have typically required complex methods, such as directed assembly of lock-and-key-shaped microgels,(12) magnetic levitation,(13) guided DNA-binding,(14) and direct 3D bioprinting of multi-ink tissue constructs(15) to name a few. To illustrate the ease of fabrication of the LEGO scaffold constructs with heterogeneous composition, a hydrogel 3D printing approach was utilized to fabricate microgels (<500 µm width and length) in the shape of cube, spiral, triangle, square, cylindrical, star, and 5-pointed flower-like geometries (**Supplementary Figure 2A, B**)(16) composed of methacrylated gelatin (GelMA), which were printed using a DLP 3D printing set up (Ember, Autodesk). Microgels were fabricated either supplemented with different combinations of human recombinant growth factors (vascular endothelial growth factor, VEGF; platelet-derived growth factor, PDGF; bone morphogenetic protein-2, BMP2); or with fluorescently-labeled microparticles (green, orange and pink), for easier visualization of spatial control (Figure 2A, B, E, F). These microgels could easily be injected in a wet or dried condition (**Supplementary Figure 2C**) or manually loaded into individual cage modules, due to their small size.

**Figure 2.**
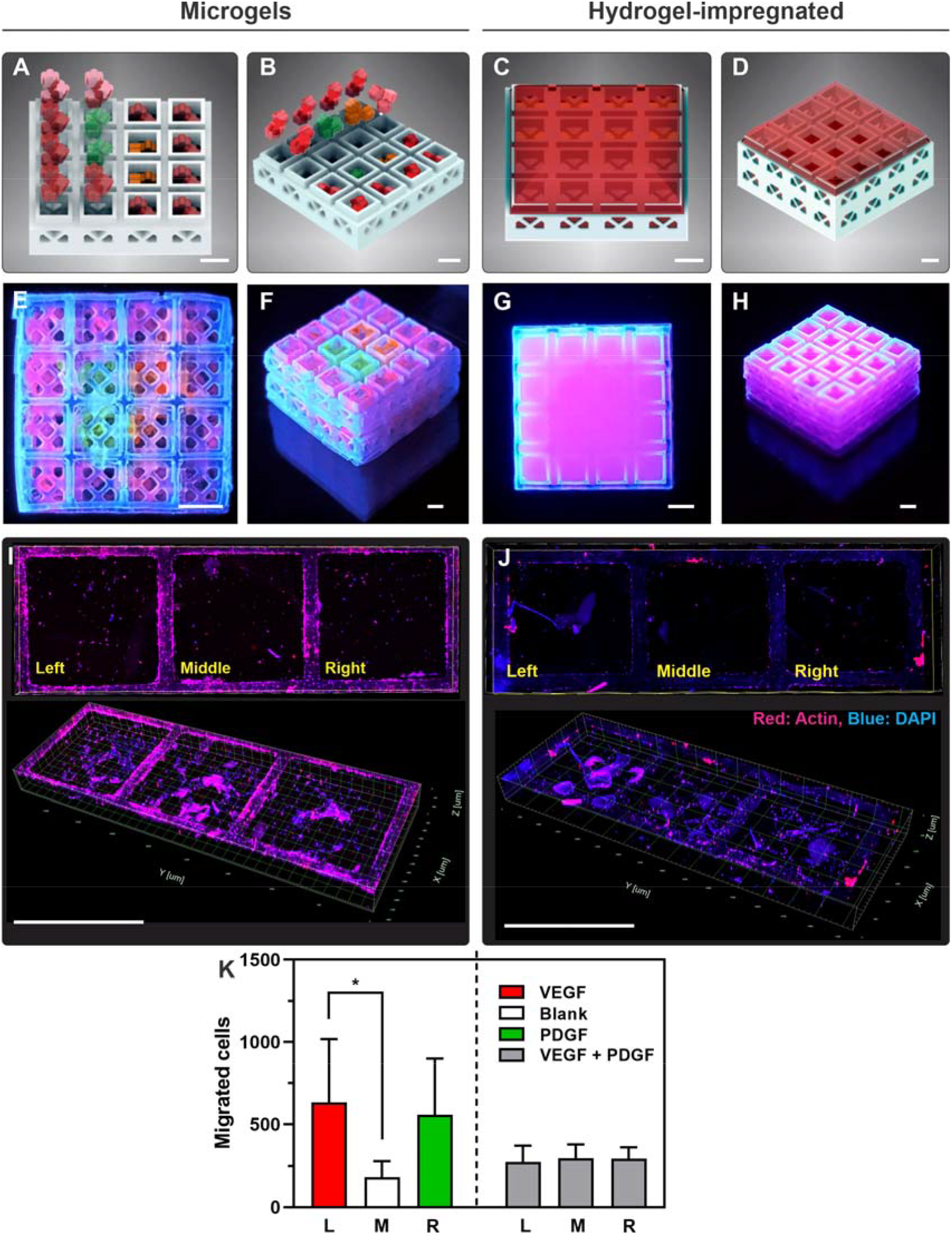
3D diagram showing the distribution of microgels in individual cage units (A, B) and the complete impregnation of GelMA hydrogel covering the LEGO scaffold (C, D), respectively. E-H) Photographs of the LEGO-like scaffolds schematically depicted above after loading with fluorescently labeled microparticles embedded with the microgel and monolithic hydrogel, respectively. I, J) Immunofluorescence staining images (Top: 2D view, Bottom: 3D view) showing directed cell migration by the microgels (loaded with VEGF [left], no growth factor [middle], PDGF [right]), when compared with the non-specific cell migration achieved by the hydrogel-impregnated group. K) Corresponding quantitative data of cells migration (p<0.05). (scale bar: 1.5 mm)

The ability to load different cages with cargo of different compositions enables the formation of multitypic constructs of various complexities with ease after individual stacks are assembled in 3D. Figure 2A, B, E, F illustrate the design and fabrication of different configurations of the 3D printed LEGO-like cages loaded with microgels encapsulated with fluorescent microparticles, distributed in different arrangements. Such a functionality is not possible when using standard hydrogel-impregnated scaffolds (*i.e.*, 3D printed, electrospun), which has long been the standard strategy for the fabrication of hydrogel-impregnated composite scaffolds in the field.(17–19) The comparison of the site-specific loading of cage modules versus a hydrogel-impregnated scaffold is shown in Figure 2A-H.

When cells were seeded in monolayer in a transwell plate positioned above engineered constructs with spatially defined presentation of VEGF and PDGF, they presented a pattern of migration that followed the chemotactic gradient of the presented cues (Figure 2I-K). VEGF was particularly attractive to the cells and stimulated significantly more cell migration than cages loaded with non-supplemented microgels (Figure 2I and 2K) (*p*<0.05). Microgels loaded with PDGF also showed directed migration, although not significantly. LEGO-like constructs fully impregnated with hydrogels carrying a mixture of VEGF and PDGF forming a composite monolith, on the other hand, showed no pattern of instructed cell migration (Figure 2J and 2K). This highlights the capability of the proposed system to function as a three-dimensionally instructive scaffold for tissue engineering. Moreover, the analysis along with the quantification suggests that when cells were able to migrate in an instructed fashion, the total cell number also appeared higher in microgel-loaded scaffolds than with hydrogel-impregnated scaffolds (Figure 2I - K), which is another advantage of the instructed approach.

Since the ultimate goal of regenerative scaffolds is to enhance the *de-novo* tissue formation, we also hypothesized that this combination of microgels and 3D printed LEGO-like cage scaffolds would enhance the diffusivity of medium carrying oxygen and nutrients into the core of a construct, in comparison to conventional scaffolds fully impregnated with a hydrogel. To test that, LEGO-like microcages were either loaded with microgels encapsulated with green fluorescence microparticles or were completely impregnated with a hydrogel prepolymer of the same composition, and then were photopolymerized to form a monolithic composite. Accordingly, the hollow characteristic of the 3D printed cages combined with the macroporosity created by the spaces between adjacent microgels enhanced the diffusion of medium, and presumably of cells and nutrients into the core of the construct (**Supplementary Figure 3A - D**), which is consistent with the patterns of advantageous diffusion of other granular hydrogels over monolithic hydrogel-based biomaterials.(20)

**Figure 3.**
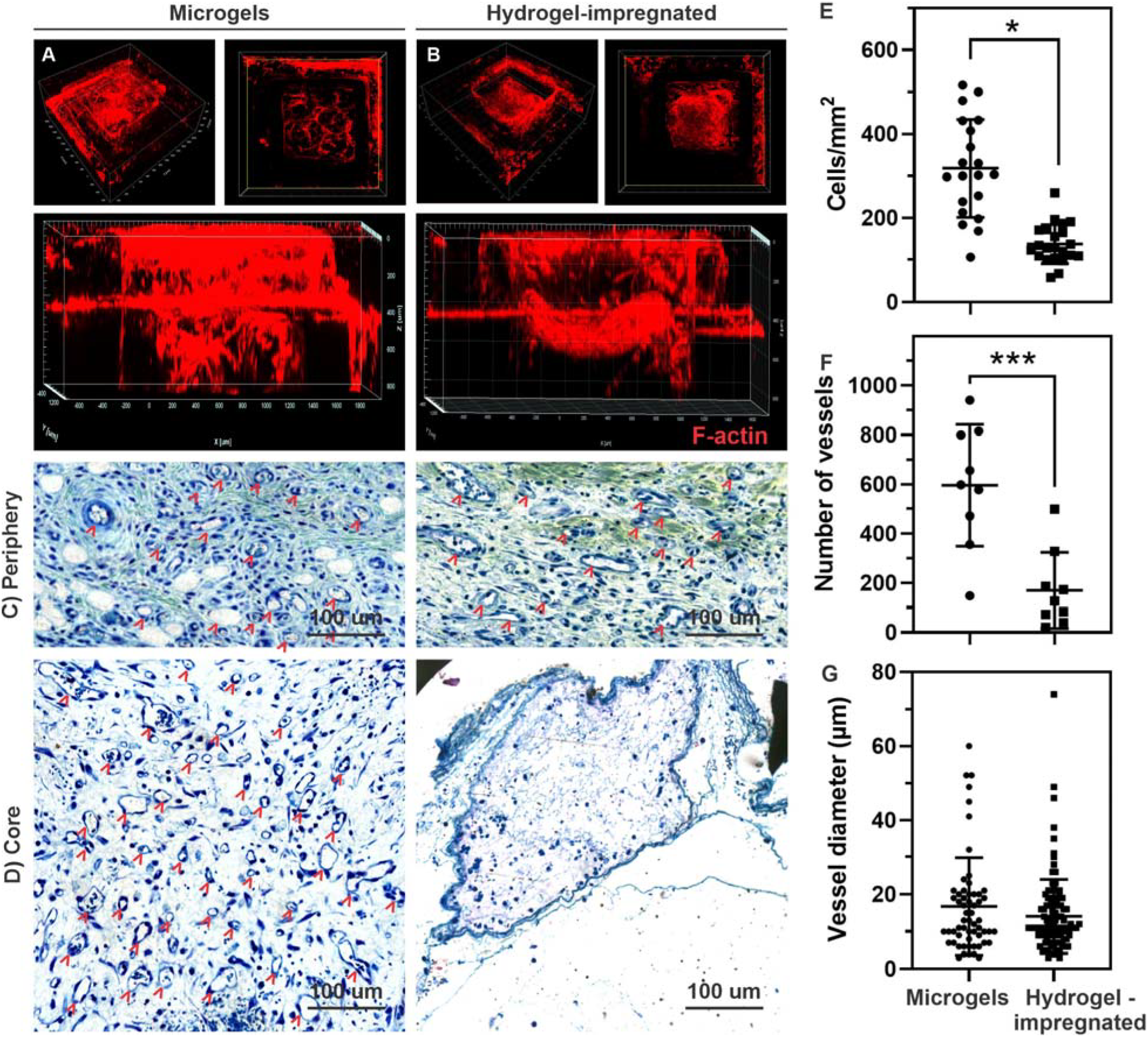
Angled, top, and orthogonal projections of F-actin stained samples show the infiltration of cells throughout the microgel-loaded LEGO-like scaffolds (A) in comparison with the hydrogel-impregnated samples (B), wherein cells are mostly found on the surface of scaffolds. C, D) Histology (toluidine blue staining) images show directed cell migration and neovascularization (indicated by arrowheads) in the periphery and core regions of the microgel-loaded LEGO-like scaffold, when compared with the hydrogel impregnated samples, which showed little evidence of cellularity and vasculature formation in the core regions. E) Quantification of cell distribution in the LEGO-like scaffolds shows nearly a 4-fold increase in the number of cells per scaffold in the microgel-loaded LEGOs in comparison to the fully impregnated scaffolds (p<0.05). F) The number of vessels and (G) average vessel diameter, respectively show a 3-fold increase in vasculature (p<0.001) and comparable diameter of formed vessels between microgel and hydrogel-impregnated LEGO scaffolds.

Subsequently, we characterized the ability of cells to invade the depth of the microgel-laden LEGO-like cages both *in-vitro and in-vivo*. LEGO-like cages loaded with microgels *in-vitro* had cells evenly distributed across the thickness of the sample, with visible actin-filament spreading throughout the construct (Figure 3A), whereas the hydrogel-impregnated cages mainly formed a monolayer of cells that were restricted to the outer surface of the construct (Figure 3B). Of note, cells appeared to conform to the outline of the microgels, thus illustrating their ability to form cell-cell and cell-matrix communication, both which appear to be primarily dictated by the pores formed between the microgels.

To compare the ability of the microgel-laden LEGO-like scaffolds to stimulate cell migration toward the core of the engineered constructs *in-vivo*, 2 stacks of 4×4 LEGO samples were implanted subcutaneously in a rat model. The hydrogel-impregnated LEGO-like scaffolds had a combination of BMP2 and VEGF homogenously mixed in the hydrogel, while microgel-filled scaffolds had VEGF-supplemented microgels in core cages, while the outer cages had microgels supplemented with BMP2. The rationale was that VEGF-loaded microgels would generate a gradient of vasculogenic factors at the core of the sample, while the pore space between the microgels would allow host vascular cells to migrate to the scaffold core within the 7 day time period, which our *in vitro* results suggested would be difficult to achieve with the fully impregnated scaffold (Figure 2). As early as one week, we detected significantly more cells distributed across the LEGO-like scaffolds loaded with microgels than in the scaffolds fully impregnated with the hydrogel as a monolith (Figure 3C, D and **Supplementary Figure 4**). The total number of cells/mm^2^ inside of the microgel-loaded LEGO scaffolds was approximately 4 times higher than that on the samples that were fully impregnated (*p*<0.05) (Figure 3E). Moreover, there was an average 3-fold increase in the total number of vessels (**Figure F**) in microgel-loaded scaffolds in comparison to the fully impregnated samples (*p*<0.01). These observations point to a significant enhancement in performance in comparison to standard hydrogel systems,(21) which have traditionally been used as monolithic grafts.(11, 22) These findings are also consistent with recent observations of implanted granular hydrogels that also demonstrate rapid healing,(20) even without the confinement of miniaturized cages in spatially controlled manner, which is a key feature of our proposed scaffold assembly method.

In summary, our group has introduced the concept of microgel-loaded LEGO-like modular material scaffolds which results in a significant increase in the ability to direct cell migration toward desirable niches with at least four times greater cell infiltration and three times higher vascularization than standard hydrogel-impregnated composite scaffolds. This represents a significant leap in performance of regenerative scaffolds by simply taking advantage of a straightforward LEGO-like design.

## Materials and Methods

### LEGO-like scaffold printing and assembly

The LEGO-like scaffold designs were generated using Computer Aided Design (CAD) software (Fusion 360, Autodesk) and 3D printed either using a previously described Lithography-based Ceramic Photopolymerization method (CeraFab 7500, Lithoz),(9) or a standard DLP 3D printer (Ember, Autodesk). In brief, similar to conventional DLP 3D printing, the LCM process is based on the selective curing of LithaBone TCP 300 (a photocurable beta tricalcium phosphate ceramic suspension) by a dynamic mask exposure process. The printing of individual layers starts with the rotation of the vat to apply a fresh slurry of a defined thickness, followed by printing, debinding, and sintering of the scaffold material. Polar resin number 48 (PR48, Ember, Autodesk) LEGO scaffolds were fabricated using standard DLP 3D printer (Ember, Autodesk), as recommended by the manufacturer. Each individual module had a hollow internal space with length, width and height measuring 1.5 mm (3.375 mm^3^), and walls measuring 560 µm in thickness. Each module formed a cage-like structure with an open top, while the walls were perforated with hollow features of approximately 530 µm - 640 µm in size separated by 200 µm – 300 µm thick struts. The top part of each module had annular protrusion of 560 µm in height, by 230 µm in thickness which interlocked precisely with a recessed feature bordered by 600 µm tall wall, also 260 µm thick (Figure 1A, 1B). Scaffolds were fabricated with repeating modules of 1×1, 2×2, 2×3 and 4×4 units, as described above. For the LEGO-like scaffold assembly process, samples were simply positioned above one another manually and stored in the required tissue culture conditions. For implantation, the assembled LEGOs loaded with hydrogels were further secured with a degradable suture line as to prevent the stacks from detaching from one another.

### Synthesis, fabrication and loading of hydrogels

Methacrylated gelatin (GelMA) was synthesized following previously published protocols.(23, 24) GelMA prepolymer at a concentration of 7-9% (w/v) was dissolved in DPBS at 80°C with 0.075% (w/v) lithium phenyl(2,4,6-trimethyl benzoyl)phosphinate (LAP, Tokyo Chemical Industry, L0290) photoinitiator. For LEGO-like scaffold impregnation with hydrogels, the 3D printed ceramics were fully immersed in GelMA prepolymer and exposed to a blue light source (405±5 nm), with a power of 20 mW/cm^2^ for 25 seconds. 3D printing of the granular GelMA microgels was also performed via a DLP printing protocol. To that end, the hydrogel prepolymer was dispensed between two polydimethylsiloxane (PDMS) (Sylgard 184, Dow Corning) sheets, separated by 600-900 μm spacers and exposed to the blue light source of the DLP printer (Ember, Autodesk) to match the wavelength, power and time specifications described above. Microgels were bioprinted in the shape of cube, spiral, triangle, square, cylindrical, star, and 5-pointed flower-like geometries, as to increase the macroporosity between adjacent gels upon scaffolds assembly (**Supplementary Figure S2**). For all experiments, the microscale hydrogel group had 5 microgels (5-pointed flower-like geometries) stacked inside each microcage, while for the hydrogel-impregnated group, the scaffolds were filled with 8-10 µl of the hydrogel prepolymer to fully fill the cage structure and surrounding walls, thus forming a monolithic composite.

### Cell culture

Human umbilical vein endothelial cells (HUVECs) were obtained from Lonza (cat #: C2519A, Basel, Switzerland). Cells were cultured in a supplemented (EGM-2 bullet kit, LONZA) endothelial cell growth medium (Lifeline cell technology, California, USA). Cell media was changed every other day and the cells were passaged when reaching a confluency of 80-90%. HUVECs from passage 4-6 were used. Primary cultures of bone marrow human mesenchymal stem cells (hMSCs) (donated by Dr. Brian Johnstone, OHSU Orthopedics) were cultured in low-glucose Dulbecco’s modified Eagle’s medium (DMEM) (Gibco, Carlsbad, USA) with 10% FBS and 1% penicillin/streptomycin. Culture media was changed every two days with one cell passage per week. hMSCs were used from passage 2-4. All cells were maintained in a humidified incubator (5% CO_2_, 37 °C).

### Cell migration and growth factor chemotaxis

To investigate cell migration and chemotaxis *in vitro*, the cell instructive LEGO-like scaffolds were prepared by loading microgels supplemented with 10 µg/mL of rhVEGF_165_ (Invitrogen, ThermoFisher) and PDGF-BB (R&D Systems) on the left and right cages of 3×1 LEGO scaffolds, respectively, while the middle cages were loaded with unsupplemented microgels (n=3). Control samples were prepared by impregnating similar 3×1 scaffolds with GelMA prepolymer loaded with similar concentrations of the growth factors, which were evenly mixed with the hydrogel material prior to loading and photopolymerization, as described above. For spatially-defined chemotaxis experiments, a co-culture of HUVECs:hMSCs in a 1:1 ratio was suspended on a density of 2 × 10^6^ cells per sample on a transwell membrane, positioned above the LEGO-like scaffold material, and allowed to migrate for a period of at least 5 days. For studies of cell infiltration into the depth of the constructs, cells were loaded directly onto the scaffolds without a membrane in a density of 2 × 10^6^ cells per well, as to mimic the immediate coverage of scaffolds with cells upon implantation and contact with blood and cell-rich extracellular fluids. After 5 days, samples were fixed with 4% paraformaldehyde (w/v) in PBS for 1 h and stained with Actin Red 555 (Molecular Probes, ThermoFisher) for 1h, rinsed with PBS and incubated with NucBlue (Molecular Probes, ThermoFisher) for 30 min at 37 °C. Scaffolds were imaged using a confocal microscope (Zeiss, LSM 880, Germany) with an objective of 10x (Zeiss, Plan-Apochromat). The depth of imaging was 100-900 µm, split into at least 250 Z-stacks. Three-dimensional (XYZ) Z-stacks were converted into TIFF files using Zen or Imaris software (v9.1, Bitplane – Oxford Instruments, Zurich, Switzerland).

### In vivo

5-7 weeks old Wistar rats were used following protocols approved by the institutional animal research ethics committee. Samples were prepared by loading the middle cages of 4×4 LEGO scaffolds with VEGF supplemented GelMA microgels and the supplementary peripheral cages with BMP2 (Shenandoah Biotechnology, Inc.) supplemented with GelMA microgels. Two stacks of scaffolds were assembled and secured with resorbable suture string prior to implantation. Control samples were prepared by impregnating similar stacks of 4×4 scaffolds with BMP2-VEGF supplemented GelMA-hydrogel and samples were implanted into four separate subcutaneous sites on the back of each rat, for a total of 6 samples per group. One-week post implantation, the scaffolds were removed, fixed in 10% formalin, embedded in paraffin and sectioned prior to staining with toluidine blue. The whole histologic slides were digitized using a Zeiss AxioScan Z1 Slide scanner at 20 X objective. Subsequently, imageJ was utilized to quantify the total number of cells inside 6 fields of each scaffold by using a color deconvolution plug-in, followed by threshold setting and automated quantification of the internal area of the scaffold. Similarly, the number and average diameter of the blood vessels was quantified using Image J.

### Statistical analysis

Statistical analyses were performed using a GraphPad Prism software 8 using a one-way analysis of variance (ANOVA) with Tukey’s multiple comparison post hoc test with an alpha (α) of 0.05.

## Acknowledgment

This project was supported by funding from the National Institute of Dental and Craniofacial Research (R01DE026170 and 3R01DE026170-03S1 to LEB), the Oregon Clinical & Translational Research Institute (OCTRI) - Biomedical Innovation Program (BIP), the Innovation in Oral Care Awards sponsored by GlaxoSmithKline (GSK), International Association for Dental Research (IADR), the Michigan-Pittsburgh-Wyss Resource Center – Regenerative Medicine Resource Center (MPW-RM), the OHSU Fellowship for Diversity and Inclusion in Research (OHSU-OFDIR to CMF). The authors declare no potential conflicts of interest with respect to the authorship and/or publication of this article.

