## Supplementary information for "3D printing of Microgel-loaded Modular LEGO-like Cages as Instructive Scaffolds for Tissue Engineering"

S. Horsophonphong

School of Dentistry, Mahidol University, Bangkok, Thailand

Dr. P. G. Coelho, Dr. Lukasz Witek

Division of Biomaterials and Biomimetics,

School of Dentistry, New York University, New York, NY, USA.

Prof. R. E. Guldberg

Phil and Penny Knight Campus for Accelerating Scientific Impact,

University of Oregon, Eugene, OR, USA

J. M. Jones, Dr. H. Xie, Dr. L. E. Bertassoni

Center for Regenerative Medicine,

School of Medicine, Oregon Health and Science University, Portland, OR, USA.

Dr. L. E. Bertassoni

Department of Biomedical Engineering, School of Medicine, and Cancer Early Detection Advanced Research Center (CEDAR), Knight Cancer Institute,

Oregon Health and Science University, Portland, OR, USA.

**Supplementary information**

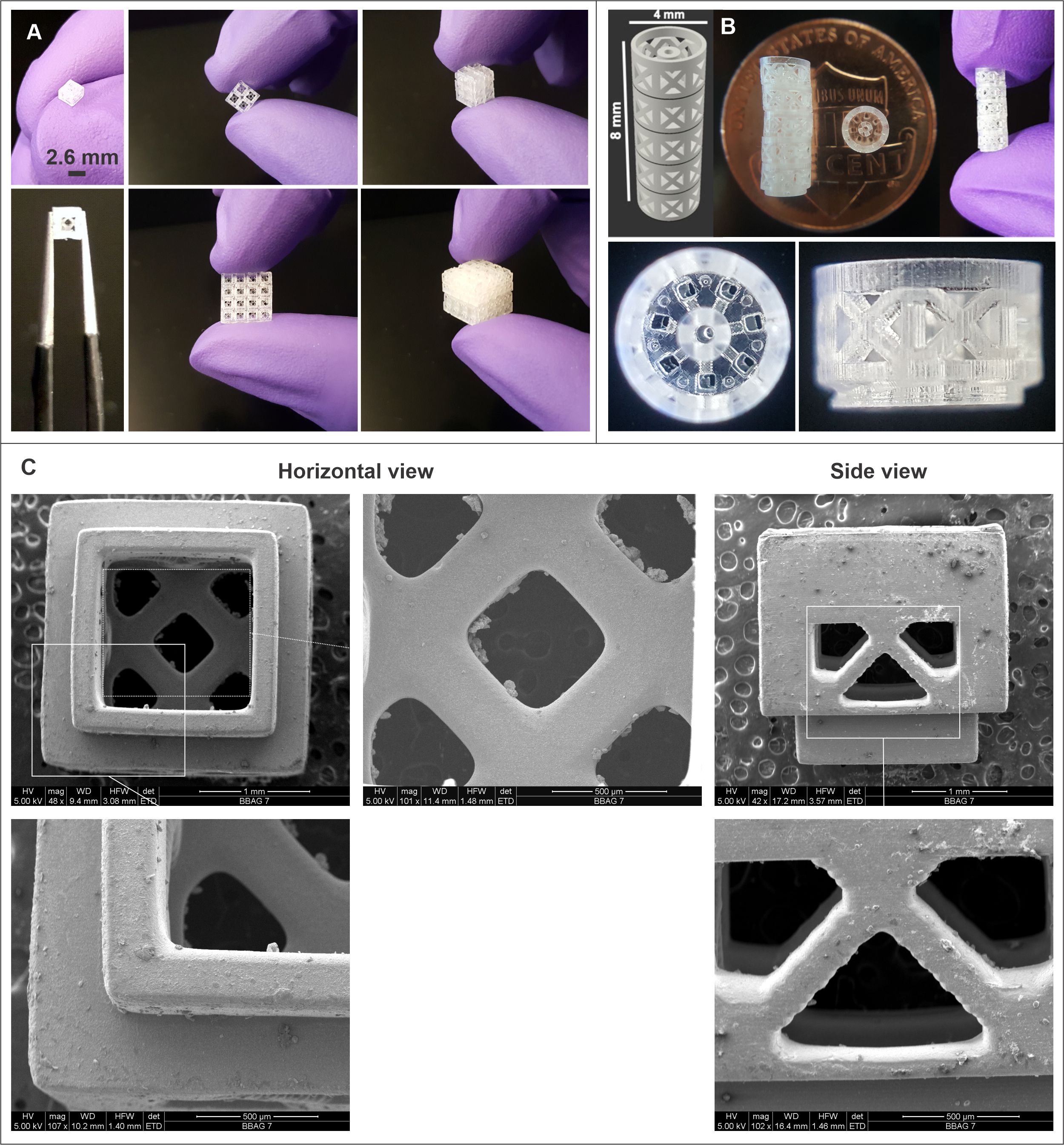

Supplementary Figure 1. A) 3D Printed scalable LEGO-like cages using PR48 resin and B) LEGO-like cylinders. C) SEM images of TCP LEGO-like scaffold from a horizontal and side view. Concentric rings and perforations on the walls allow for significantly higher cell invasion when used with microgels.

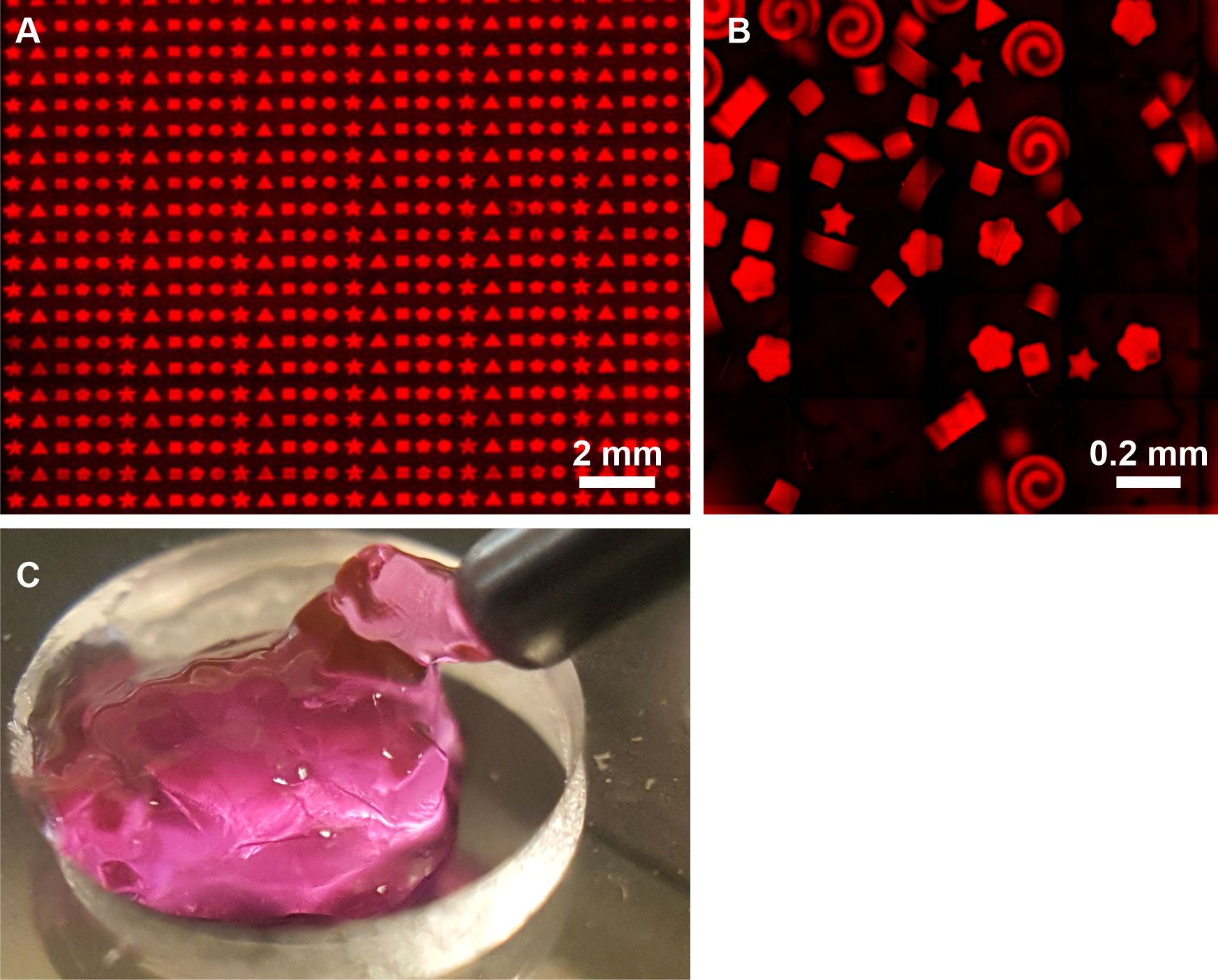

Supplementary Figure 2. Print capability of high throughput hydrogel microarrays. A) Array of DLP-printed microgels with over 1K constructs in the shape of triangle, square, pentagon, star, and five-pointed flower-like geometries, B) ready-to-dispense DLP printed microgels, and C) dispensing of freshly prepared microgels through a dispensing needle.

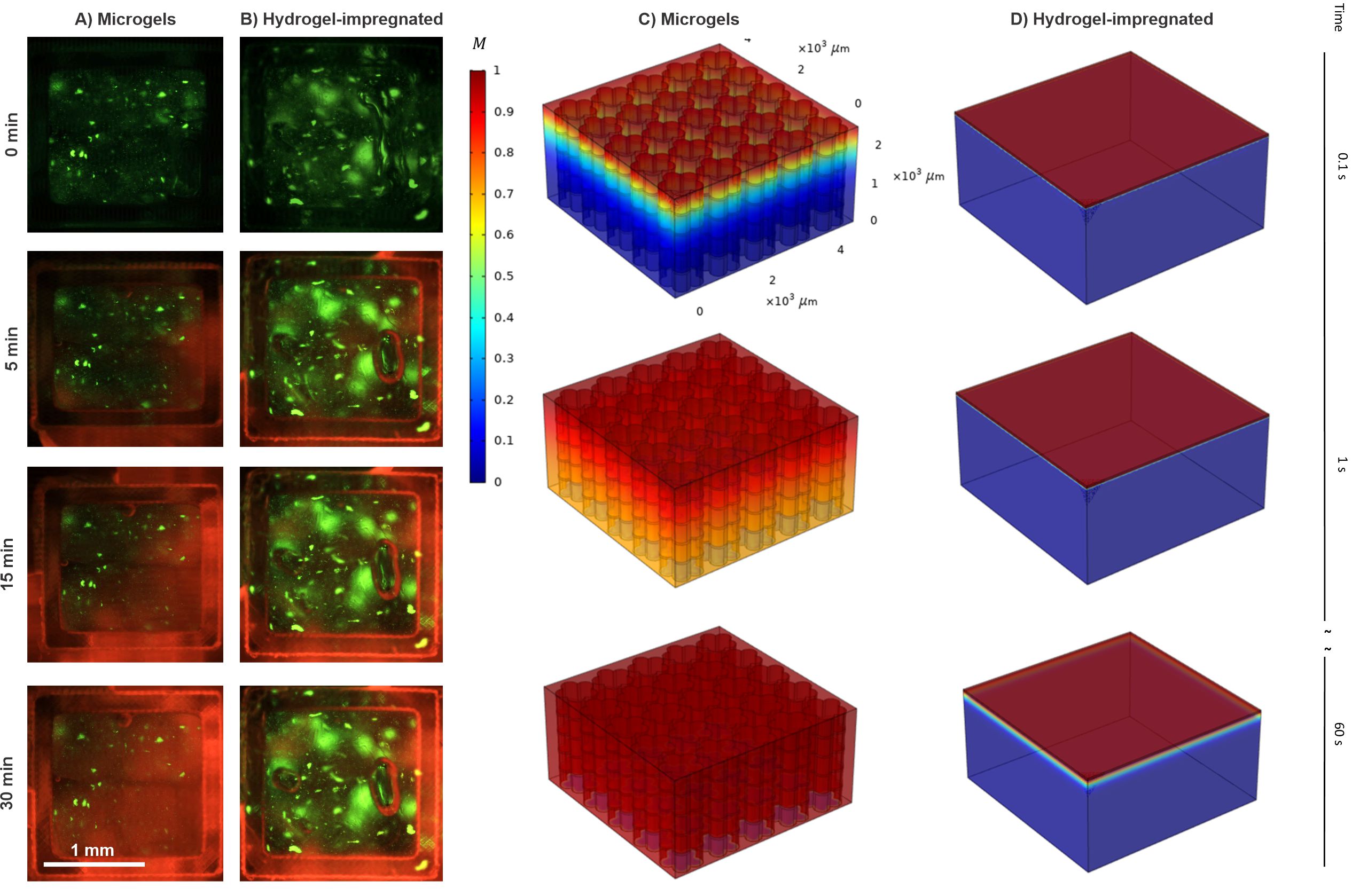
Supplementary Figure 3. Rhodamine diffuses from top to bottom region in a faster rate when the LEGO scaffold is loaded with A) microgels than B) hydrogel-impregnated. In order to evaluate the diffusion behavior of biomolecules through microgels, the species diffusion is formulated by the diffusion equation, $\frac{\partial c}{\partial t}+\nabla\cdot\left( -D\nabla c \right)=0$, where c and D represent species concentration and diffusion coefficient. The diffusion coefficient of Rhodamine B inside the gel and solution is set as 2.47e^-6^ and 4.50e^-6^ cm^2^/s, respectively. In both cases, the top boundary is set as C=1 M, and all the other outer boundaries have no flux condition. The diffusion behavior of Rhodamine B in packed flower-shaped microgels (C), and hydrogel-impregnated (D) at 0.1, 1, and 60 s. Results clearly demonstrate the rapid diffusion of the solution through the microgels than the hydrogel-impregnated.

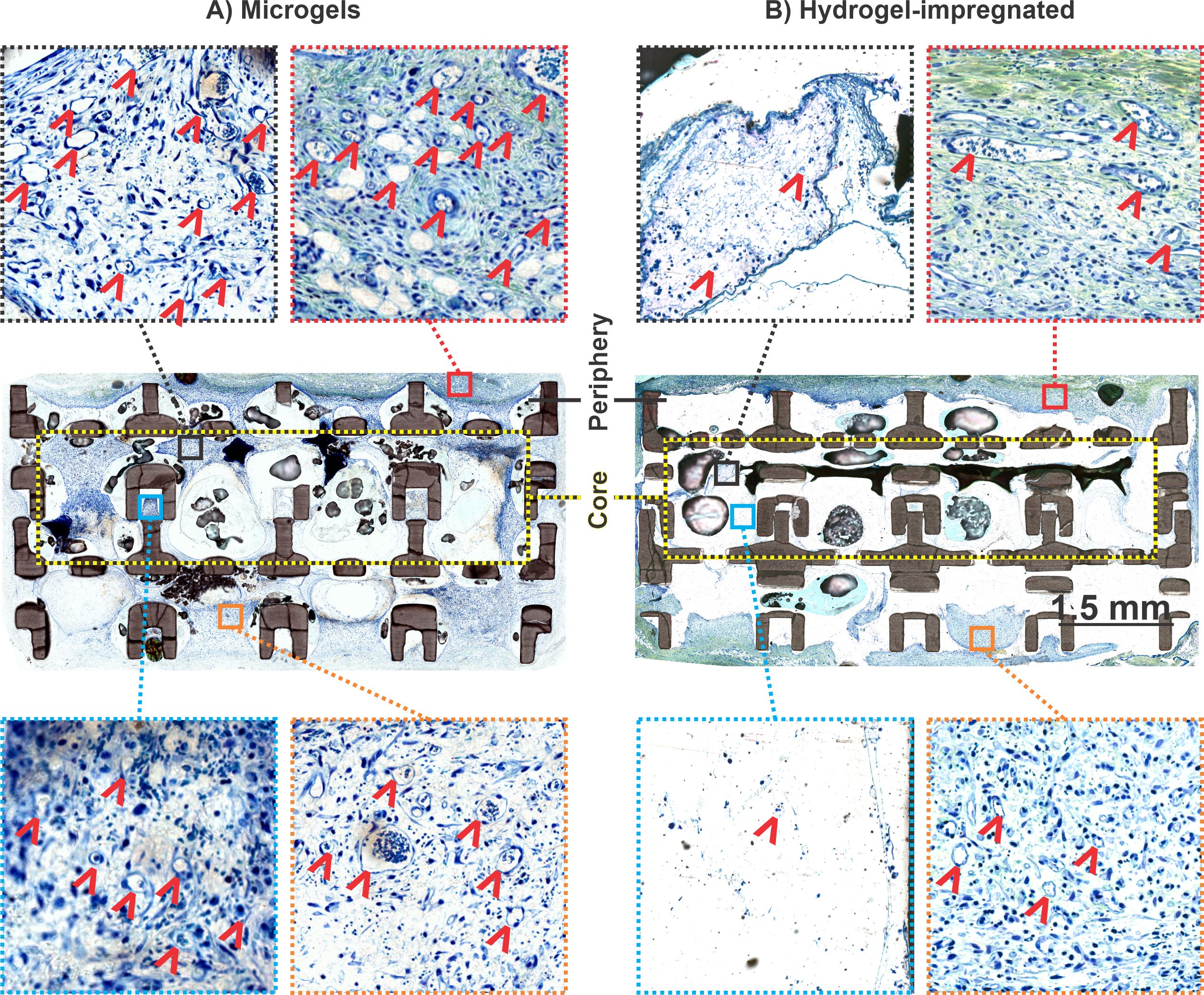

Supplementary Figure 4. Histology (Toluidine blue staining) images of the whole LEGO-like scaffold loaded with microgels (A) and hydrogel impregnated (B), respectively. High magnification images taken at the core (black and blue squares) and periphery regions (red and orange squares), show increased migration of cells and neovascularization (indicated by arrowheads) in the periphery and core regions of the microgels loaded LEGO-like scaffold.
